# Diversity, composition, and networking of saliva microbiota distinguish the severity of COVID-19 episodes as revealed by an analysis of 16S rRNA variable V1-V3 regions sequences

**DOI:** 10.1101/2022.10.20.513136

**Authors:** Violeta Larios, Beatriz Meza, Carolina Gonzalez, Francisco J Gaytan, Joaquín González Ibarra, Clara Esperanza Santacruz Tinoco, Yu-Mei Anguiano Hernández, Bernardo Martínez Miguel, Allison Cázarez Cortazar, Brenda Sarquiz Martínez, Julio Elias Alvarado Yaah, Antonina Reyna Mendoza Pérez, Juan José Palma Herrera, Leticia Margarita García Soto, Adriana Inés Chávez Rojas, Guillermo Bravo Mateos, Gabriel Samano Marquez, Concepción Grajales Muñiz, Javier Torres

**Author notes:** Corresponding author: Javier Torres.

## Abstract

**Background:** Studies on the role of the oral microbiome in SARS-CoV-2 infection and severity of the disease are limited. We aimed to characterize the bacterial communities present in the saliva of patients with varied COVID-19 severity to learn if there are differences in the characteristics of the microbiome among the clinical groups.

**Methods:** We included asymptomatic subjects with no previous COVID-19 infection or vaccination; patients with mild respiratory symptoms, positive or negative for SARS-CoV-2 infection; patients that required hospitalization because of severe COVID-19 with oxygen saturation below 92%, and fatal cases of COVID-19. Saliva samples collected before any treatment were tested for SARS-CoV-2 by PCR. Oral microbiota in saliva was studied by amplification and sequencing of the V1-V3 variable regions of 16S gene using a Illumina MiSeq platform.

**Results:** We found significant changes in diversity, composition, and networking in saliva microbiota of patients with COVID-19, as well as patterns associated with severity of disease. The presence or abundance of several commensal species and opportunistic pathogens were associated with each clinical stage. Patterns of networking were also found associated with severity of disease: a highly regulated bacterial community (normonetting) was found in healthy people whereas poorly regulated populations (disnetting) were characteristic of severe cases.

**Conclusions:** Characterization of microbiota in saliva may offer important clues in the pathogenesis of COVID-19 and may also identify potential markers for prognosis in the severity of the disease.

**Importance of the work:** SARS-CoV-2 infection is the most severe pandemic of humankind in the last hundred years. The outcome of the infection ranges from asymptomatic or mild to severe and even fatal cases, but reasons for this remain unknown. Microbes normally colonizing the respiratory tract form communities that may mitigate the transmission, symptoms, and severity of viral infections, but very little is known on the role of these microbial communities in the severity of COVID-19. We aimed to characterize the bacterial communities in saliva of patients with different severity of COVID-19 disease, from mild to fatal cases. Our results revealed clear differences in the composition and in the nature of interactions (networking) of the bacterial species present in the different clinical groups and show community-patterns associated with disease severity. Characterization of the microbial communities in saliva may offer important clues to learn ways COVID-19 patients may suffer from different disease severities.

## Introduction

SARS-CoV-2 infection is the most severe human pandemic in the last one hundred years, with over 599 million cases and 6.46 million deaths worldwide as of August 2022. By 2.5 years after it first appeared, the virus has become endemic, with occasional outbreaks that have sometimes required re-implementation of public health containment measures. SARS-CoV-2 infects human cells expressing the angiotensin-converting enzyme 2 (ACE2), which is used as receptor for the spike protein. This property enables the virus to infect different cells in different organs, causing a multisystem, multiorgan infection (1,2). The outcome of infection ranges from asymptomatic or mild episodes to severe and fatal cases (3). Severity of disease is known to be determined by multiple factors, including host and viral characteristics.

Another factor that may determine severity of disease is the human microbiome, which is known to play a major role in the modulation of other infections. A healthy microbial community may mitigate the transmission, symptoms and severity of viral infections (4). Members of our normal bacterial communities may also inhibit viral replication and modulate the inflammatory response to counteract the infection and prevent immune mediated tissue damage (4). However, studies on the role of the microbiome in SARS-CoV-2 infection and on severity of the disease have been limited. All studies reported to date have found that patients infected with SARS-CoV-2 have an altered microbiome composition in nasopharyngeal samples, with reduced diversity, although differences in bacterial composition vary among studies. One study reported that Propionibacteriaceae were significantly more abundant and *Corynebacterium accolens* significantly decreased in infected patients (5). Others reported that *Prevotella* and *Alloprevotella* were increased in abundance in severe cases (6). Metatranscriptomic analyses of nasopharyngeal swabs and sputum samples also found a reduced diversity in patients with COVID-19 pneumonia when compared with non-COVID-19 pneumonia. Different species of *Prevotella, Veillonella, Haemophilus, Fusobacterium* and *Gemella* showed a reduced abundance in the COVID-19 cases (7). The oral mucosa has been recognized as an important site for SARS-CoV-2 infection and as a source for spreading the infection to the upper and lower respiratory tract (8). It has been reported that the oral microbiome (tongue swabs) forms a dysbiotic pattern in long-COVID cases, with higher abundance of microbiota that induce inflammation, including *Prevotella* and *Veillonella* (9). Diversity of oral (tongue scraping) microbiome was also found reduced in patients with COVID-19, with increases in bacteria producing lipopolysaccharide and decreases in bacteria producing butyric acid (10).

The oral cavity hosts over 1,000 bacterial species, representing the second largest and most diverse microbial community in the human body after the gut (11). The oral microbiota has been shaped as a result of coevolution with the human mouth. Local niches in the mouth supply shelter and nutrients, whereas the microorganism complement the physiological needs of the host, including nutrients, immune training, and host defense (12). The oral microbiota is formed by a collection of compositionally distinct communities that reflect the array of diverse microenvironments present in the different regions of the mouth. These communities usually grow as highly structured symbiotic biofilms, linked through physical and metabolic associations that confer a fitness advantage to the entire microbial community and make them particularly stable and resilient to microenvironmental changes (13). The salivary microbiota has been shown to be a conglomerate of bacteria shed from oral surfaces in the pharynx, the tongue and the tonsils as the main sites of origin (14). Thus, saliva is an appropriate sample that mirrors the. microbiota from different regions of the oral cavity. Saliva can also be aspirated and reach the lungs, representing an important source of infection for the respiratory tract. Indeed, studies have suggested a relationship between oral hygiene and respiratory diseases, including asthma, chronic obstructive pulmonary disease, and pneumonia (15).

In the present work we aimed to characterize the bacterial communities present in the saliva of patients with COVID-19. The study included groups of patients with different severity of disease, from mild to fatal cases, in order to determnie if there were differences in the composition of the microbiome among clinical groups. The results demonstrated clear differences in diversity, composition and networking of the bacterial species present in the saliva of the different clinical groups.

## Material and Methods

### Patients studied

The study was done during the period of June 2020 to January 2021 a period of high epidemiological uncertainty and strict containment measures because of COVID-19. Therefore, recruitment of patients was done by the attending health personnel. Under these circumstances we did not have enough information to estimate a sample size, nor were we certain of the feasibility to reach an expected number of cases. Thus, patients were invited to the study as they were presented with suspicious symptoms, and before a confirmed diagnosis in mild cases, or within five days of hospitalization in severe confirmed cases. Final sample size of the clinical groups was based on the inclusion-exclusion criteria.

Asymptomatic cases (AC) were individuals without symptoms, no previous COVID-19 infection (as referred by the patient) or SARS-CoV-2 vaccine, and no antibiotics in the last four weeks. Smokers and those with any chronic disease were excluded. Mild cases were ambulatory patients with mild respiratory symptoms (fever, cough, headache, odynophagia, myalgias) presenting for COVID-19 diagnosis. They were sampled before any treatment, including antibiotics, and followed until recovery. After testing for SARS-CoV-2 by PCR test (COBAS 6800, Merck México, Mexico city), they were classified as positive (AP) or negative (AN) for the infection. Patients who during follow up required hospitalization because they developed severe symptoms were excluded from this group and included in the hospitalized group. Severe patients were cases that required hospitalization because of severe symptoms (HP), particularly an oxygen saturation below 92% and the presence of comorbidities of risk including hypertension, diabetes, morbid obesity, immunocompromise, cardiovascular or neurological diseases, chronic renal failure, tuberculosis or neoplasia. Patients were usually hospitalized within the first seven days after symptoms started and followed until discharged because of recovery or improvement (treated at home until recovery) or because of death. For the analyses, patients that died were included in a group of deceased cases (DHP).

### Saliva samples

A volume of 10 ml of saline solution (0.85% NaCl) was given to patients, who were asked to thoroughly wash the mouth and spit back into a 50 ml plastic tube. Samples were immediately transported to a central laboratory for SARS-CoV-2 diagnosis using a PCR test to amplify a fraction of the spike and N protein genes. On arrival samples were immediately inactivated by heating at 65ºC for 30 min. After the diagnosis, samples were sent to our laboratory for microbiome studies. Transport of the samples was done following international regulations for safety, including special multiple packing with dry ice. Once received in our lab, samples were frozen to -70ºC until studied.

### DNA extraction

Saliva samples (1 mL) were centrifuged for 10 min at 5000 x g, then bacterial pellets were suspended and incubated at 37°C for 3 hours with 180 μL of enzyme solution (20 mg/mL lysozyme; 20 Mm Tris-HCl, pH 8.0; 2 mM EDTA; Triton® 1.2%). Subsequently, DNA was extracted using the QIAamp® DNA Mini Kit (Qiagen) according to the manufacturer’s protocol.

### Preparation of the amplicon libraries from the 16S rRNA hypervariable regions V1-V3 and sequencing

The amplicons of the V1-V3 hypervariable regions of the 16S rRNA gene were generated using previously published primers (16), which were ligated to the adapter sequences (17) and used to assemble the DNA libraries. For library assembly, 25 ng of DNA were mixed with 12.5 ul of Go-Taq green master mix enzyme (Promega) and 10 uM of each primer (supplementary table 1), and amplified using the following conditions: 3 min at 98° followed by 25 cycles (20 sec at 98°C, 30 sec at 65°C, 30 sec at 70°C), 5 min at 72°C and 4°C hold. Libraries were normalized and sequenced using the 2×251 cycle configuration, with 20% Phix control, and 100 uM of the sequencing primers (see Supplementary Table 2) and placed in positions 12, 13 and 14 of the sequencing cartridge (2). Sequencing of the libraries was performed on the Miseq platform (Illumina, San Diego CA, USA).

### Bioinformatics analysis

### Quality control and taxonomic classification

Paired sequencing fastq files were QC inspected for phred values and for absence of adapters using FastQC v0.11.9 (18). The data were processed using the DADA2 v1.20.0 pipeline (19). Standard filter parameters (maxN = 0, truncQ = 15, and maxEE = 2, lengths below 200 bp were discarded) were used. The process of reads continued with dereplication filtering, and removal of chimera formation (representing < 2%) using the removeBimeraDenovo option. 16S sequences associated with chloroplast or mitochondria were removed. The sequences were grouped into Amplicon Sequence Variants (ASVs) with the naive RDP Bayesian classifier of DADA2, and taxonomic classification was assigned to the species level using the Human Oral Microbiome Database (20) (eHOMD 16S V1-V3 training set).

### Alpha and beta diversity

The alpha and beta diversity were determined using the Phyloseq (21), Vegan (22), and ggplot2 (23) packages to obtain the Chao1 (richness estimator), Fisher (abundance estimator), and the Shannon and Simpson (diversity and evenness estimators) values. The ape package v5.3 (24) was used for the generation of the phylogenetic tree. *Statistical significance* was evaluated with a univariate ANOVA analysis and p-values ≤ 0.05 were considered significant. Between-group differences in beta diversity were assessed using non-metric multidimensional scaling (NMDS) with the Bray-Curtis dissimilarity index to visualize the clustering of the individual samples. Significance was assessed by calculating non-parametric permutational multivariate analysis of variance (PERMANOVA) with 10,000 permutations, using the adonis and beta disper functions. Analysis of similarities (ANOSIM) (25) and homogeneity of group dispersions (PERMDISP) were also performed.

### Biomarker discovery

The differences in taxa between the experimental groups were studied in order to identify the ASVs that significantly distinguished each group (marker bacteria). The analyses were done using different models, including Random Forest (RF) with 1,000 trees to build the model and LEfSe (14) with a qvalue ≤ 0.1 and linear discriminant analysis (LDA score ≥ 2.0) (26). To analyze the differences in taxa abundance among groups, the Fold Change (FC) and the FDR-adjusted were calculated using DESeq2 (27), and volcano plots were constructed with EnhancedVolcano v1.12.0 (28).

### Composition of the core microbiome

The microbiome v1.17.3 package with the core function (29) was used to calculate central Taxa among all groups. Samples were filtered applying a prevalence of 0.5 and a detection of 20% (threshold for absence/presence) among the samples. The visualization of the data was carried out through a heatmap with the plot core function, the variables captured were relative abundance and the ASVs of the output samples for all the studied groups.

### Co-occurrence networks

A single association network was built with the NetCoMi v package (30) applying agglomeration at the species level and using the netconstruct function (measure of correlation of “pearson”, zeroMethod of “multRepl” and normMethod “clr”) and the netAnalyze function (clust_fast_greedy method). Graphics were generated using the Fruchterman-Reingold layout algorithm of igrap. The size of the nodes was adjusted by a normalization of the counts, a color was assigned to each subcluster within the net, and the single nodes were removed. Estimated associations are shown with green connections for positive or red for negative correlations.

### Metabolic routes

The inference of metabolic pathways was generated with the ASV’s sequences using Picrust2 (31) and the KEGG database (32) for both Level 2 and Level 3. The z-score of relative abundances and clustering were calculated using the package pheatmap v1.012.

## Results

The number of patients that consented to participate were close to 500 and from those we were able to have enough saliva sample and follow up clinical data in 390 cases. After excluding patients who did not fulfill inclusion criteria, including quality and amount of sample, 314 saliva samples were sequenced. In the end, 278 samples (88.5%) passed the Q30 quality value with an average of 200,000 reads per sample and a total yield of 16.1 Gb, distributed as described in Table 1.

**Table 1.** Characteristics of patients with COVID-19 studied for oral microbiome in saliva samples.

| Group of patients | Number studied | Age mean ±sSD | Sex ratio male:female |
| --- | --- | --- | --- |
| Asymptomatic | 32 | 27.8±7.7 | 0.78 |
| Mild negative* | 70 | 38.2±12.1 | 0.59 |
| Mild positive* | 102 | 41±13.1 | 0.77 |
| Severe positive** | 57 | 50.8±13.9 | 1.94 |
| Deceased** | 17 | 74.4±6.5 | 3.3 |

\*ambulatory patients; \*\*hospitalized patients

### Diversity and abundance of bacterial species differ among patients with different severity of COVID-19

Microbial diversity indexes of the saliva samples are presented in Figure 1a; results show that richness (Chao index) was higher in the asymptomatic individuals but gradually and significantly decreased in the ambulatory, hospitalized, and deceased patients, a result that was further supported by the Fisher analysis. Shannon index however showed an increased value in all symptomatic groups, suggesting an increase in evenness in these patients. Diversity in the microbial composition among groups was analyzed by a NMDS Bray analysis (figure 1b), which showed some separation of the bacterial communities in the clinical groups. These differences in bacterial composition were more evident when the abundance of species was analyzed across groups (figure 1c). By amplifying the V1-V3 regions of the 16S rRNA gene and by using the eHOM database to annotate, we were able to identify most of the AVS to the level of species (20). We identified a group of species that was abundant in asymptomatic adults but decreased in ambulatory patients and was almost absent in the hospitalized and deceased patients, including *Gemella elegans, Neisseria sicca, Porphyromonas gingivalis* and *Rothia dentocariosa*. There were also species whose abundance was higher in ambulatory patients than in either healthy or hospitalized cases, including *Actinomyces graevenitzii, Mogibacterium diversum, Streptococcus mutants* and *Actinomyces odontolyticus*. Finally, a number of species had a higher abundance in hospitalized patients, including the deceased cases, such as *Staphylococcus epidermidis, Capnocytophaga granulosa, Prevotella melaninogenica* and *Leptotrichia wadei*. Of note, the abundance of the opportunistic pathogen *Acinetobacter baumannii* was particularly higher in the deceased patients than in any other group.

**Figure 1.**
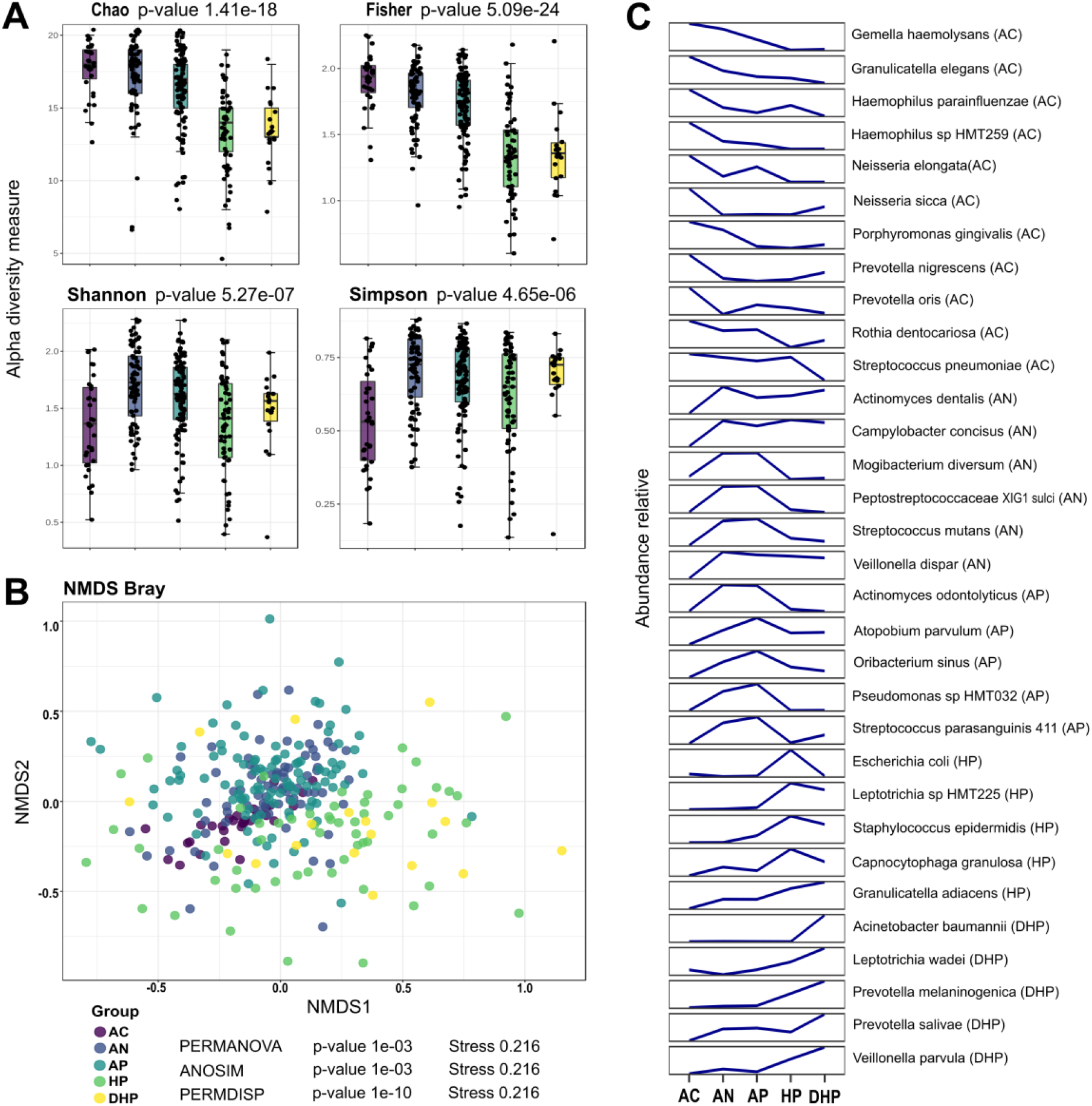
Indices of microbial diversity and relative abundance of bacterial species in saliva of patients with different severity of COVID-19 disease. A) Chao index, Fisher analyses, Shanon and NMDS Bray analyses; B) Beta diversity with the NMDS Bray-Curtis dissimilarity index; C) analyses of abundances of species across groups.

### Pairwise comparisons showed marked and significant differences in bacterial composition among groups

Pairwise differences between groups were studied with the enhanced volcano test (Figure 2). Compared with the asymptomatic group, *Actinomyces odontolyticus, Streptococcus parasanguinis, Oribacterium sinus, Atopobium parvulum* and *Streptococcus mutants* (among others) were significantly more associated with the mild ambulatory group (Figure 2a); in contrast, *Prevotella intermedia, Porphyromonas gingivalis, Alloprevotela sp*. and *Prevotella oris* were more associated with healthy adults. When the group of hospitalized patients with severe disease was compared with the healthy individuals, *Leptotrichia sp, Escherichia coli, Staphylococcus epidermidis* and *Prevotella oris* were significantly more associated with HP patients (Figure 2b). *Haemophilus sp*. HTM259, *Porphyromonas gingivalis, Actinomyces sp*. HTM169, *Haemophilus parainfluenzae* and others were more associated with the asymptomatic group. *Acinetobacter baumannii, Capnocytophaga granulosa, Prevotella melaninogenica, Granulicatella adiacens, Prevotella salivae*, and *Veillonella parvula* were significantly more associated with the DHP fatal patients compared to asymptomatic patients (Figure 2c). In contrast, *Granulicatella elegans, Haemophilus parainfluenza, Veillonella sp*. HMT780, *Alloprevotella sp*. HMT308 and *Prevotella intermedia* were more associated with asymptomatic adults. Of interest, when the two groups of hospitalized patients were contrasted, *Acinetobacter baumannii* and *Prevotella salivae* were more associated with deceased patients, whereas *Escherichia coli, Leptotrichia* HMT215 and *Staphylococcus epidermidis* with the severe HP patients (Figure 2d). Thus, *Acinetobacter baumannii* and *Prevotella salivae* were marker species that differentiated deceased patients from severe hospitalized patients and from asymptomatic individuals.

**Figure 2.**
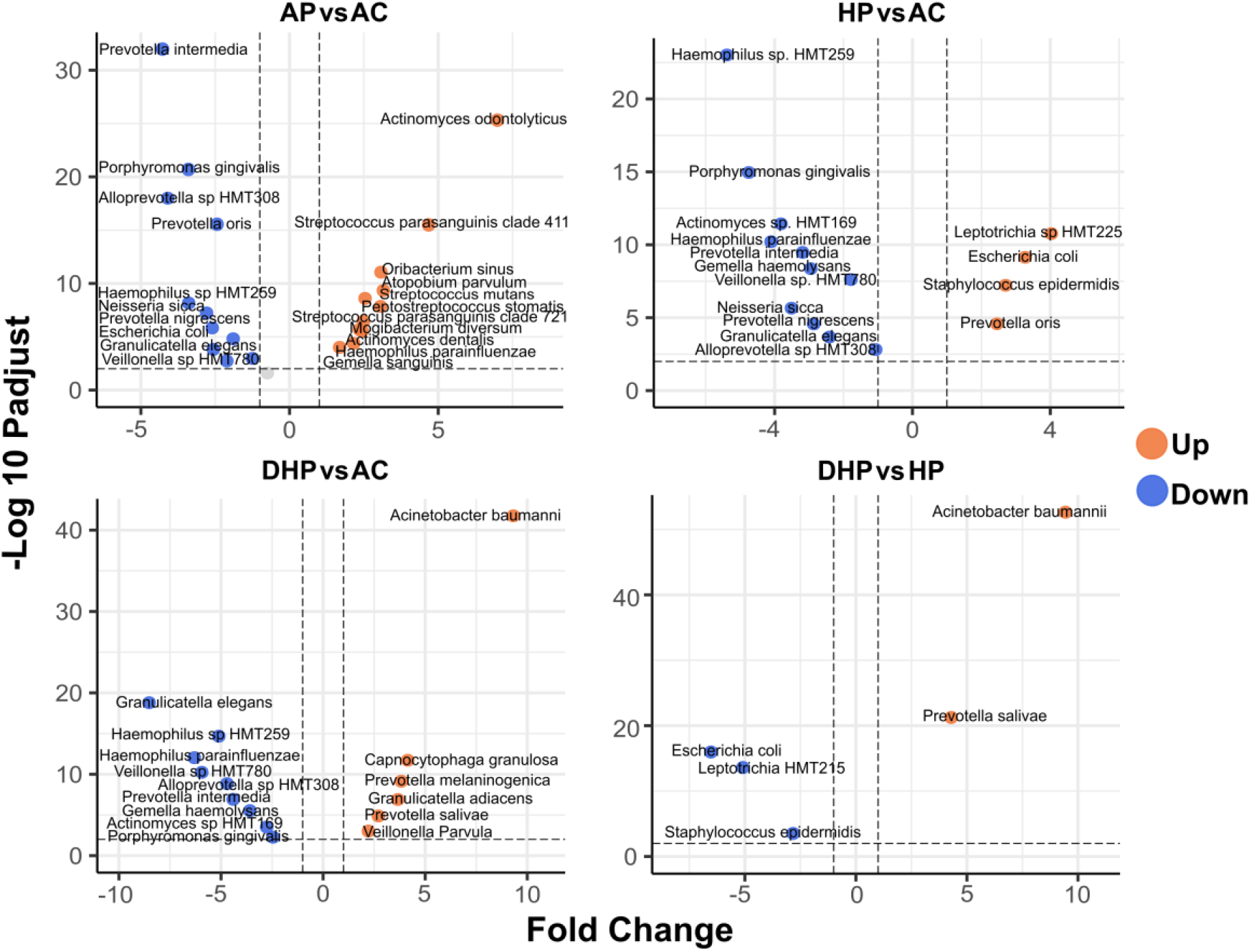
Pairwise comparison between clinical groups analyzed with the enhanced Volcano test. A) comparison between the ambulatory SARS-CoV-2 positive group and the asymptomatic control group; B) comparison between the hospitalized SARS-CoV-2 positive group and the asymptomatic control group; C) comparison between the deceased SARS-CoV-2 positive group and the asymptomatic control group; D) comparison between the deceased and the hospitalized group. Species in orange dots presented increased abundance and those in blue decreased abundance.

We also studied a group of patients with mild respiratory disease that were negative for SARS-CoV-2 infection (AN). Compared with the asymptomatic adults (suppl Fig 1a), these patients had higher abundance of *Actinomyces graevenitzii, Streptococcus mutants, Peptostreptococcacceae* XI G1, *Actinomyces dentalis* and *Stomatobaculum longum* whereas *Prevotella intermedia, Veillonella sp* HMT780, *Alloprevotella sp* HMT308, *Escherichia coli* and *Neisseria sicca* were more abundant in the healthy group. Of note, when the two groups with mild disease (AP and AN) were compared, *Streptococcus parasanguinis* was more abundant in the SARS-CoV-2 patients, whereas *Veillonella dispar, Peptostreptococcacceae* X1 G1, *Porphyromonas pasteri. Actinomyces dentalis* and *Actinomyces graevenitzii* were significantly more abundant among the mild non-infected patients (Suppl Fig 1b).

### An all-vs-all comparison revealed species that distinguishes each clinical group

We next examined the differences among all groups by means of the random forest and LEfse analyses (Figure 3). In this all-vs.-all groups’ analyses, random forest (Figure 3a) found *Haemophilus parainfluenzae, Prevotella nigrescens, Neisseria sicca, Gemella haemolysans* and *Rothia dentocariosa* among those most significantly distinguishing the asymptomatic group, whereas *Streptococcus parasanguinis, Oribacterium sinus, Atopobium parvulum* and *Actinomyces odontolyticus* differentiated the mild ambulatory cases. *Escherichia coli, Leptotrichia sp* HTM225 and *Staphylococcus epidermidis*, distinguished the hospitalized patients, whereas *Veillonella parvula, Prevotella melaninogenica, Acinetobacter baumannii, Granulicatella adiacens, Actinomyces dentalis, Capnocytophaga granulosa, Leptotrichia wadei* and *Veillonella dispar* strongly differentiated the deceased patients.

**Figure 3.**
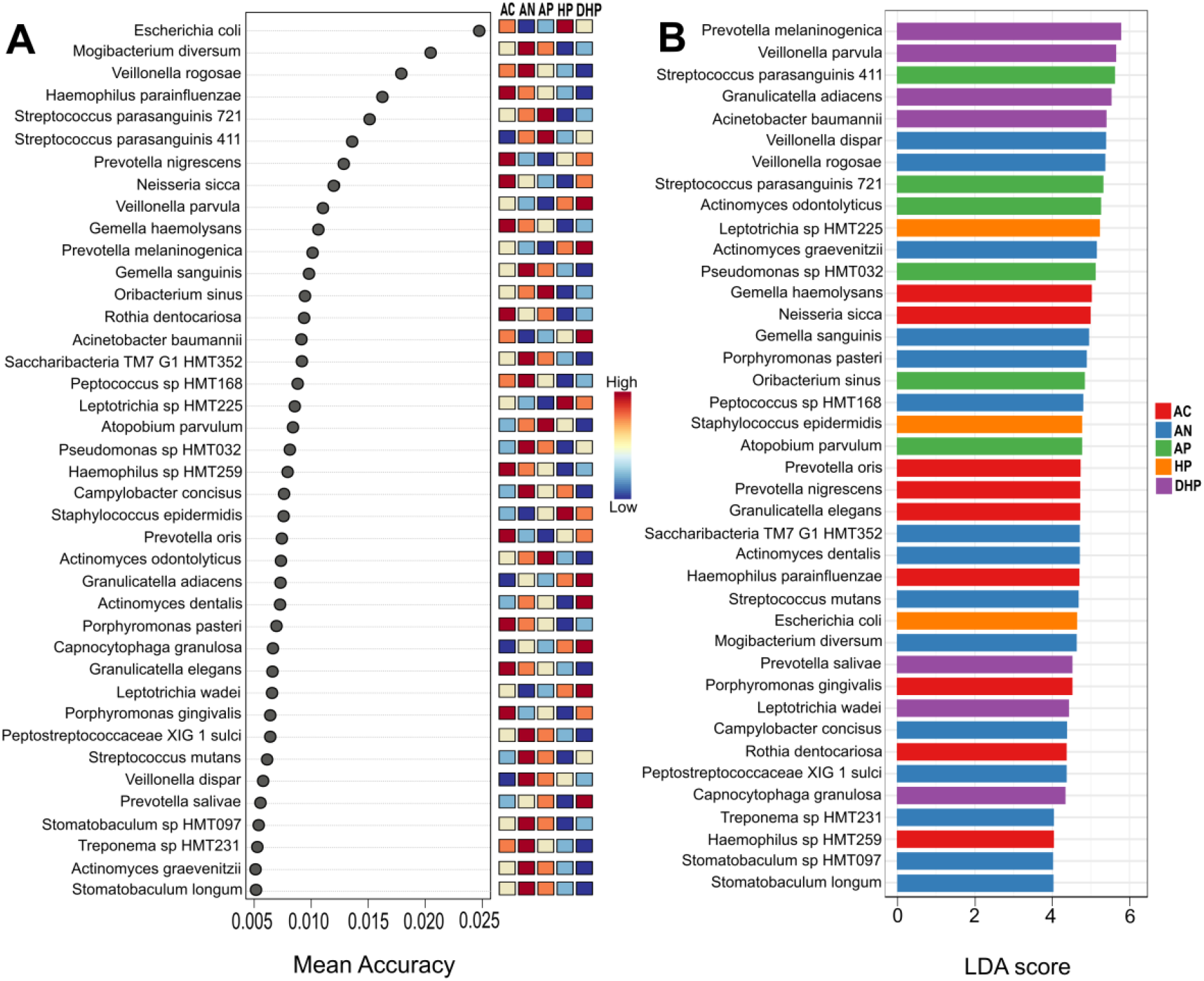
Analyses of all-vs-all clinical groups by A) Random forest; B) Lefse test. Groups are indicated to the right of the figures, CA asymptomatic, AN ambulatory SARS-CoV-2 negative, AP ambulatory SARS-CoV-2 positive, H hospitalized, HD deceased patients.

Results with the linear discriminant analysis (LDA) (Figure 3b) showed a strong agreement with random forest and of note, the two models pointed to *Prevotella melaninogenica, Veillonella parvula* and *Acinetobacter baumannii* as species strongly distinguishing the group of deceased patients, whereas *Neisseria sicca, Haemophilus parainfluenzae* and *Prevotella nigrescens* were markers for asymptomatic adults.

### A core microbiome analysis show species present in all clinical groups

Finally, we determined the core microbiome (Figure 4) to learn which species were present in all clinical groups, probably because they are more resilient to changes in the microenvironment. Of note, *Streptococcus pneumoniae* was found present in all clinical groups with a relative abundance of over 10% in over 60% of the patients (see also Suppl Figure 2), highlighting its endurance regardless of the clinical condition of the patients. *Granulicatella adiacens, Veillonella dispar, Streptococcus parasanguinis, Prevotella melaninogenica* and *Veillonella parvula* showed a relative abundance of over 1.0% in at least 50% of all patients (Suppl Figure 2). Other species of *Actinomyces, Prevotella* and *Veillonella* were also among the species in the core microbiome.

**Figure 4.**
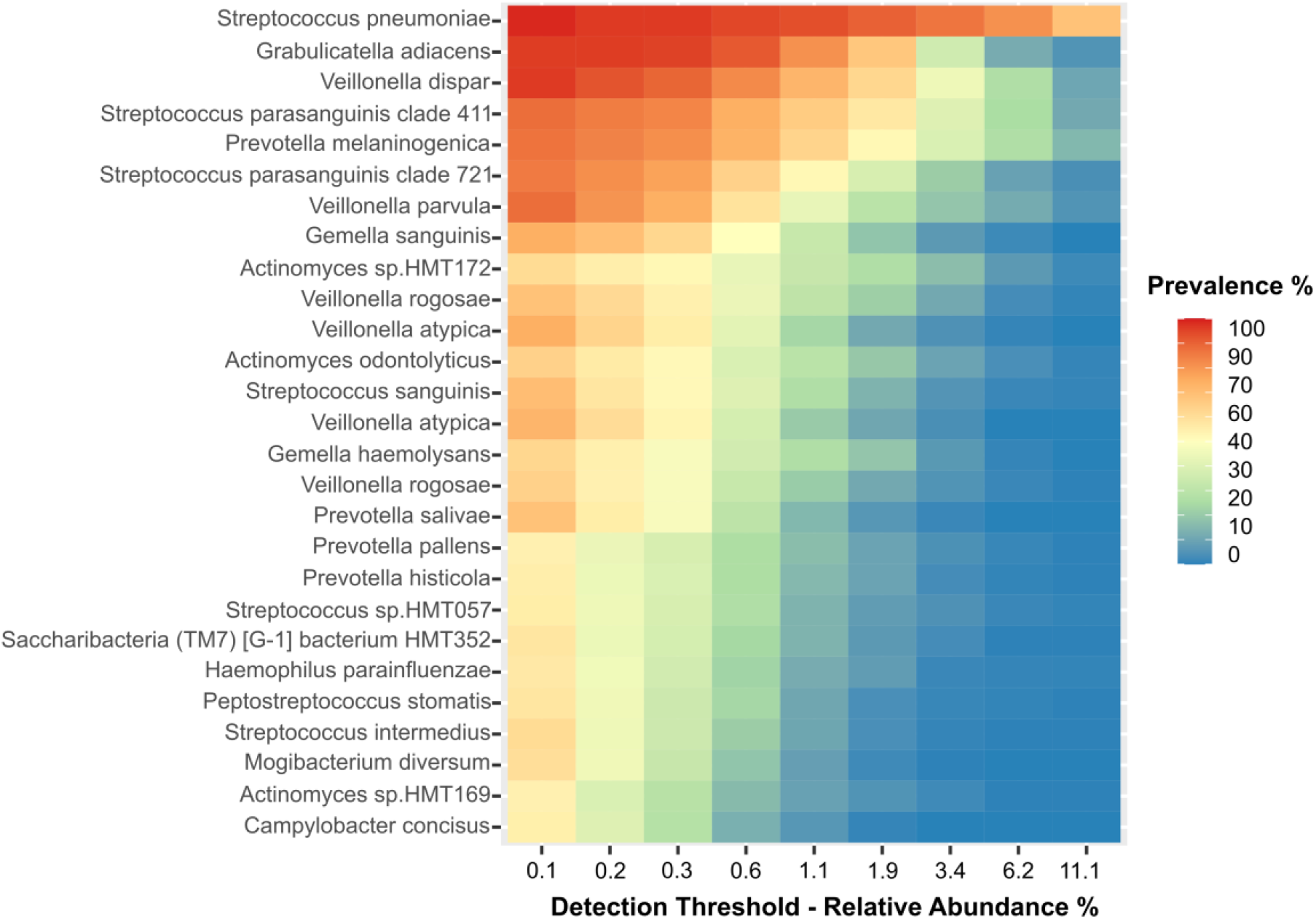
Determination of the core microbiome for all clinical groups. The relative abundance is described on the X axis and for each species a heatmap for the prevalence is shown for each relative abundance.

### Analysis of networking in bacterial communities shows marked contrasts in the different clinical groups

The interaction between the members of the bacterial community in each group was studied by network analyses. In each network, the size of the circle is proportional to the abundance of the species and each circle’s color represents subclusters of bacteria with a stronger interaction between them. The color of each connection (edge) relates to the type of interaction, green is positive, and red is negative. For these analyses, we selected a group of species based on the previous results in the Volcano, LEfse, and random forest analyses. We selected up to 50 species that showed significant association with each of the five clinical groups studied and results are presented in figure 5 and Table 2.

**Table 2.**
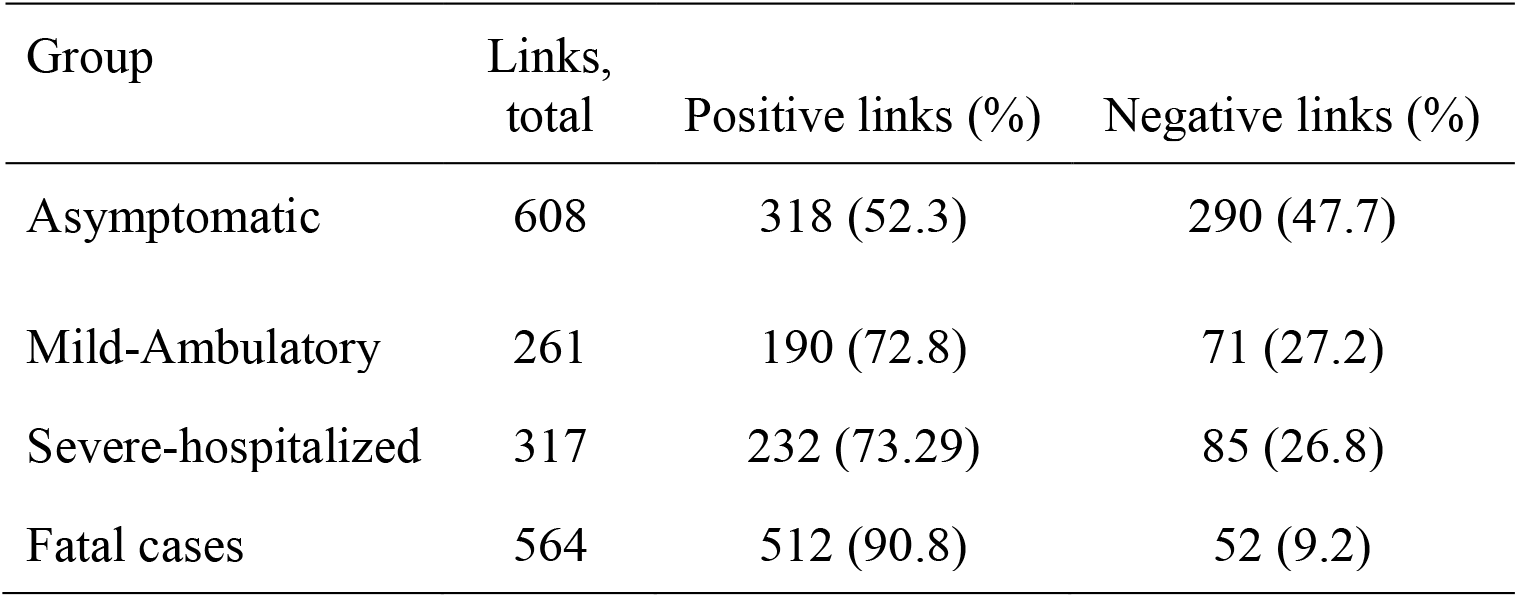
Number and type of links (edges) in the bacterial network in saliva of patients from different clinical groups.

**Figure 5.**
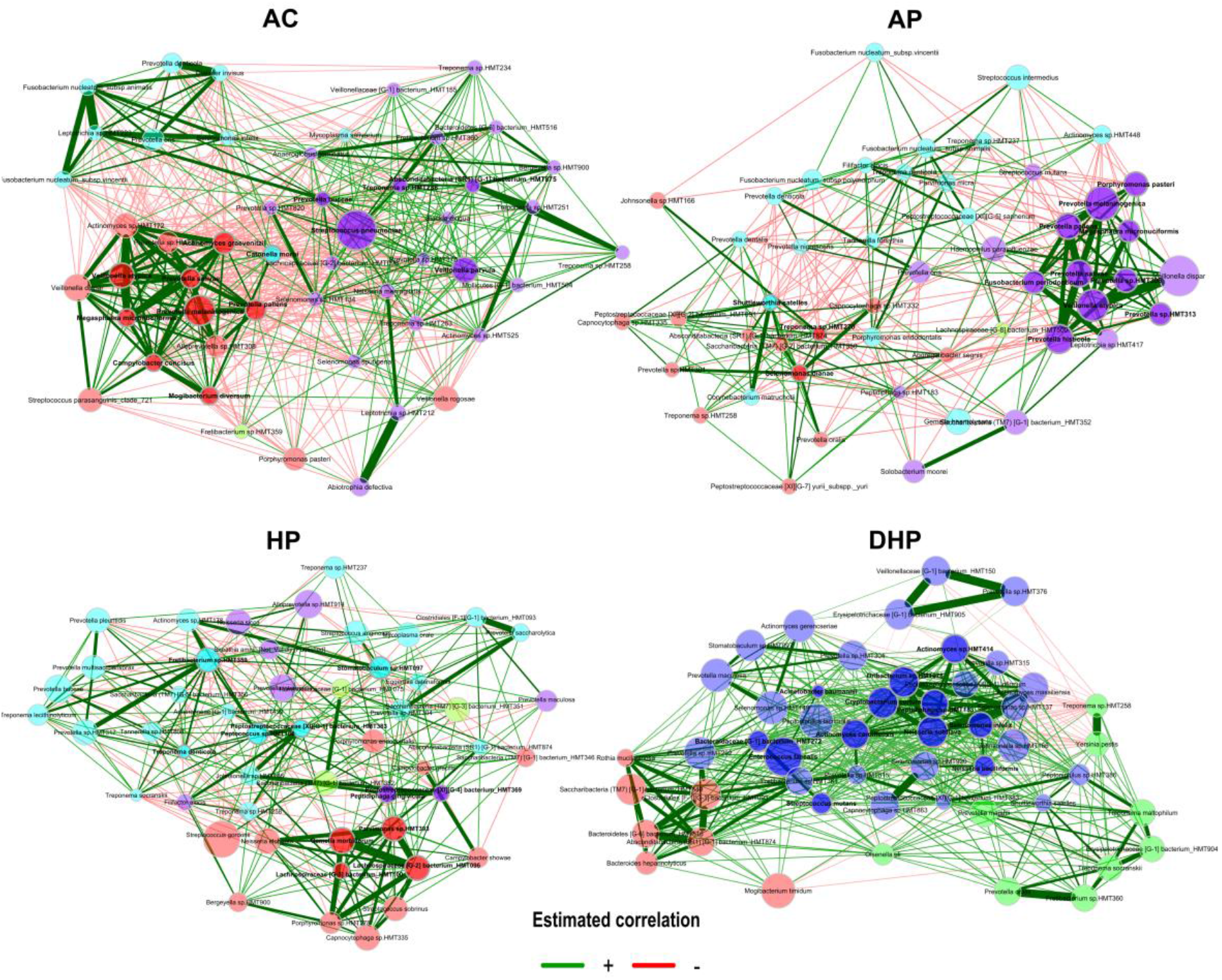
Co-occurrence networks for each clinical group analyzed by Pearson correlation with the NetCoMi v package. The size of the nodes was adjusted by a normalization of the counts, and a color was assigned to each subcluster within the network. Positive associations are shown with green connections and negative correlations with red connections.

In the asymptomatic group the net shows *Streptococcus pneumoniae* as the most abundant species, placed in the center of the network with many links with other members of the community (Figure 5a). Compared with the other groups, this population presented a higher number of edges (608), and a balance in the positive (52%) and negative (48%) correlations, showing a highly interactive and regulated community. We also observed subclusters with strong positive interaction within their members but negative interaction with members of other subclusters. One such subcluster (red-dark and red-light circles) is formed by *Prevotella salivae, Prevotella melaninogenica, Prevotella pallens, Prevotela sp*. HTM313, *Veillonella atypica, Veillonella dispar, Actynomices sp*. HT172, *Actinomyces graevenitzi, Megasphaera micronuciformis, Campylobacter consisus, Alloprevotela sp*. HTM308 and *Mogibacterium diversum*. Two other subclusters are apparent, one in light blue and the other purple-dark and purple-light.

In the ambulatory-positive group the number of subclusters and edges (261) were markedly reduced, with the positive correlations more frequent (73%) than the negatives (27%) (Figure 5b). One subcluster present in the asymptomatic patients was partially present here, with *Prevotella salivae, Prevotella melaninogenica, Prevotella pallens, Prevotella sp* HTM313, *Veillonella dispar, Veillonella atypica*, and *Megasphaera micronuciformis* as members, but with *Prevotella histicola, Leptotrichia sp*. HTM417 and *Fusobacterium periodonticum* as new elements. In this network it is still evident the presence of negative interactions with members of the other subclusters, although to a lesser extent.

Structure of the hospitalized patients’ network was clearly different, with a reduced number of interactions (317), most of them positive correlations (73%) (Figure 5c). *Fretibacterium sp*. HTM359 showed a strong positive interaction with many members of the group. This species has been found abundant in deep periodontal pockets in patients with periodontitis and has been suggested as a biomarker for periodontitis (33). In this group, *Neisseria elongata, Gemella morbittorum, Parvimonas sp* HTM393, *Lachnospiraceae* bacterium HTM100 and HTM096, *Porphyromonas sp*. HTM278, *Capnocytophaga sp*. HTM335 and *Streptococcus sobrinus* created a strongly interactive subcluster. The behavior of the community in the deceased patients was strikingly different, showing a large number of edges (564) in a densely interactive community, with the vast majority of interactions being positive (91%) and most of the few negative correlations coming from two species: *Yersinia pestis* and *Treponema sp*. HTM258. Three subclusters with strong interaction within them but also positive interaction with members of the other subclusters were apparent. One subcluster included: *Rothia mucilaginosa, Saccharibacteria* (TM7), *Bacteroides heparinolyticus, Bacteroidetes* (G-6)HMT516, *Absconditabacteria* (g-1)HMT874 and *Clostridiales* (f-1)HMT093. A second subcluster was *Actinomyces sp*. HMT414, *Actinomyces cardiffensis, Cryptobacterium curtum, Oribacterium* HMT078, *Peptidiphaga* HMT183, *Neisseria subflava, Selenomonas sp*. HMT137 and *Selenomonas infelix*. The third strong subcluster included only *Erysipelotrichaceae* HMT905, *Prevotella sp*. HMT376 and Veillonellaceae (G-1)HMT150. Of note in this community is the presence of *Acinetobacter baumannii* and *Enterococcus faecalis* both with intense interaction with other members and which were almost absent in the asymptomatic and mild cases.

### Estimation of metabolic activity shows a highly increased activity by the bacterial community of the severe and deceased patients

An approximation to the metabolic activity of the bacterial community in each group was deduced with the use of Picrust software. Notably, the analyses revealed a significantly increased metabolic activity by the bacterial community of deceased patients in most of the metabolic pathways, which gradually decreased in hospitalized patients, in ambulatory patients and finally in asymptomatic adults (Figure 6). This drastic tendency is clearly illustrated looking at the metabolic pathways at the two different levels presented in Figures 6a and 6b. The few activities diminished in severe and deceased patients were the biosynthesis of the antimicrobials aminoglycosides (produced by actinomycetes) and clavulanic acid (Streptomyces) as well as the metabolism of xenobiotics by P450 and polyketide sugar biosynthesis (Figure 6b).

**Figure 6.**
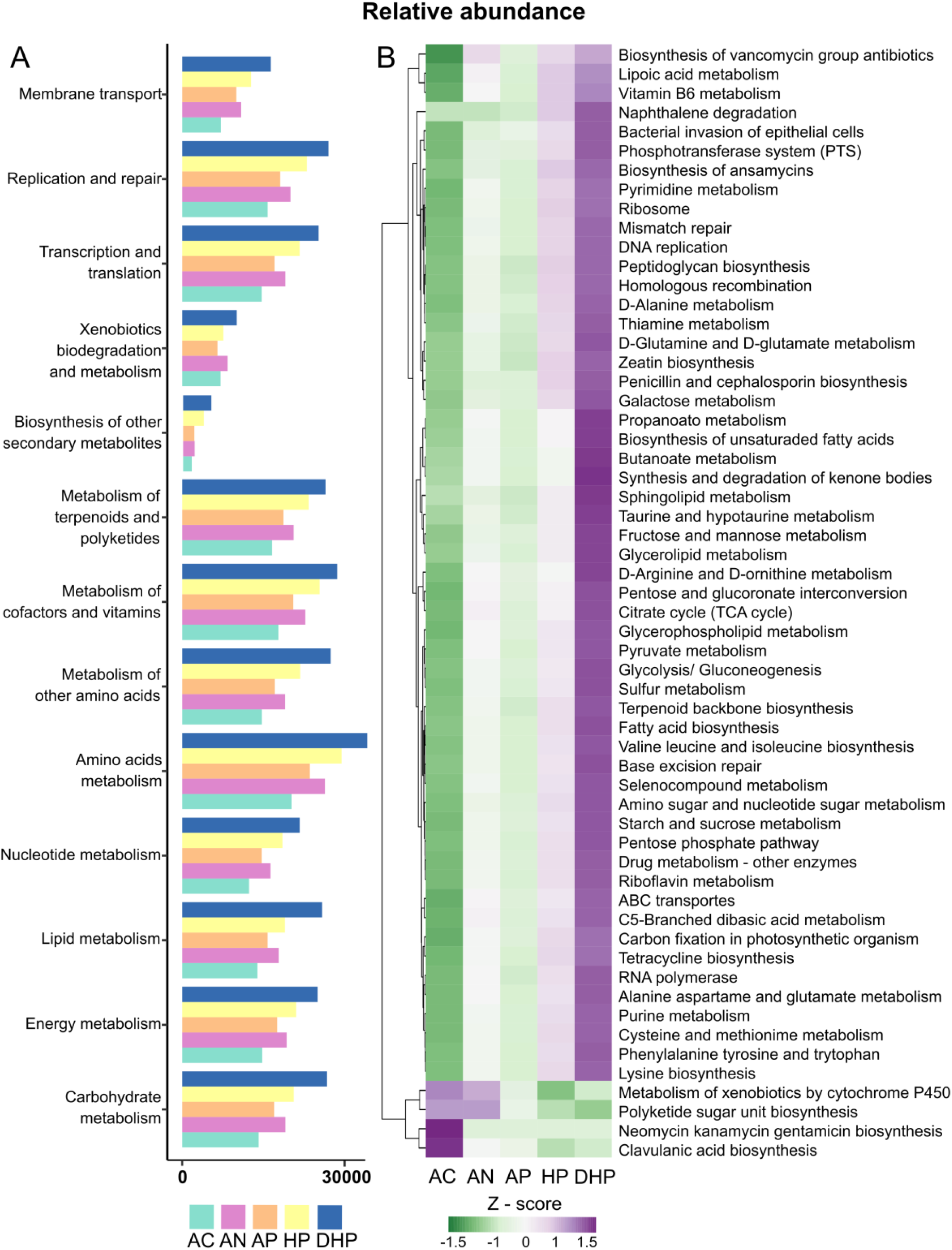
Metabolic activity by the bacterial community present in each clinical group as inferred by Picrust2 software and KEGG database. Figures present the relative abundance of the metabolic pathways level 2 (A) or level 3 (B) as defined in KEGG.

## Discussion

The oral cavity may be the entry to the respiratory tract and the source of the oral microbiome of the upper and lower airways, including the lungs. In fact, the oral mucosa is recognized as an important site for SARS-CoV-2 infection and as a source for spreading the infection to the upper and lower respiratory tract (8). Thus, it becomes relevant to study the oral microbiome in patients with COVID-19 to try to elucidate its role in the severity of the disease. In this study we used saliva as a surrogate of the oral microbiota (14).

Our results show that diversity of the bacterial communities in saliva decreases as the severity of the disease increases, from ambulatory to hospitalized to deceased patients. Previous studies reported similar results when comparing healthy controls vs COVID-19 patients (10) or vs patients with long-COVID (9), but these studies did not contrast oral microbiota in patients with different severity of disease. Several studies in nasopharyngeal samples have consistently reported reduced diversity in COVID-19 patients (5),(6,7) indicating infection is associated with important changes in the structure of bacterial populations colonizing the upper airways. To further study the nature of the changes in the microbiota we searched for differences in its composition among the groups. A main difference of our work with previous COVID reports is that we amplified the V1-V3 region of the 16S rRNA gene, which results in improved sensitivity when working with oral microbiome (34) and we used the updated eHOM database with an extended coverage of oral species (eHOM) (20). Thanks to this we were able to identify most ASV to the level of species, which contrasts with previous studies usually reporting down to the level of genus. This is relevant if we consider that the oral cavity hosts over 1,000 bacterial species (11) and reporting differences at the level of genus fell short in the interpretation of results. This can be illustrated by our findings that *Prevotella intermedia* and *Prevotella oris* were found significantly associated with asymptomatic adults, whereas *Prevotella melaninogenica* and *Prevotella salivae* were associated with deceased cases. These differences would be missed if analysis is limited to the genus level.

Volcano test, random forest and LEfse analyses showed a strong agreement in severity-associated species and led us to identify specific changes in the composition of bacterial communities that differ in patients according to the severity of the disease. Thus, our results show that *Gemella haemolysans, Neisseria sicca, Prevotella oris, Prevotella nigrescens, Rothia dentocariosa, Porphyromonas gingivalis*, and *Granulicatella elegans* are salivary markers of asymptomatic adults. Previous studies have also reported *Granulicatella elegans* more prevalent in healthy subjects. *Porphyromonas gingivalis* is associated with periodontal disease and is considered a “keystone pathogen” because of its ability to modify the host inflammatory response and cause dysbiosis (35). *Streptococcus parasanguinis, Actinomyces odontolyticus, Oribacterium sinus* and *Atopobium parvulum* were characteristic of mild COVID cases, *Leptotrichia sp* HMT225, *Staphylococcus epidermidis* and *Escherichia coli* of severe cases, and *Prevotella melaninogenica, Veillonella parvula, Granulicatela adiacens, Acinetobacter baumannii, Prevotella salivae, Leptotrichia wadei* and *Capnocytophaga granulosa* of fatal cases. A comparison with previous studies is difficult because most of them report up to genus level, but also because studies with oral microbiota are scarce (9) (10). Even studies with nasopharyngeal samples are limited and they show contradictory results. Whereas some agree reporting that *Corynebacterium* significantly decreased and *Prevotella* increased in infected patients (5),(6),(36), others have reported a reduction of *Prevotella* and *Veillonella* in COVID cases (7,35).

Changes in the composition of bacterial communities are not the only factor to consider in microbiota studies. Coinfections with pathogens or opportunistic pathogens are common during viral pneumonia and known to increase the severity and risk of mortality (37). Thus, *Streptococcus pneumoniae* coinfection is a major cause of increase morbidity and mortality during influenza infection (38). Coinfections have also been documented in patients with COVID-19, particularly with *S. pneumoniae, K. pneumoniae* or *H. influenza* (39), and a metagenomic study of nasopharyngeal samples of patients with COVID-19 found a co-infection with a clinically relevant microorganism in 12.5% of patients (5). It should be noted that in our work *Streptococcus pneumoniae* was present in all groups studied and in fact its abundance was higher in healthy individuals and decreased as the severity of the disease increased, which questions its role as an opportunistic pathogen in our population.

In our study coinfections were common in severe hospitalized cases: *Escherichia coli, Leptotrichia* HMT215 and *Staphylococcus epidermidis* in severe patients and *Acinetobacter baumannii* and *Leptotricia wadei* in fatal cases. *Leptotrichia* has been found increased in patients with COVID-19 (10). *Leptotrichia* species are present in the oral cavity of healthy individuals and is considered an opportunistic pathogen because its abundance increases in caries, stomatitis or cases of septicemia in immunocompromised patients. *L. wadei* has been isolated in saliva of patients with caries or halitosis (40). On the other hand, *Acinetobacter baumannii*, a hospital-acquired opportunistic pathogen, has been reported in severe COVID patients, even in the lungs of fatal cases (41) and we found it significantly more abundant in the saliva of deceased patients.

We also determined the core microbiome for all clinical groups, which is seldom reported in microbiome studies. As indicated above, *Streptococcus pneumoniae* was found with high relative abundance in over 80% of all cases, representing the most prevalent and most abundant species in our population. The next most prevalent species included *Granulicatella adiacens, Veillonella dispar, Streptococcus parasanguinis* and *Prevotella melaninogenica*, with a relative abundance of over 1.0% in most cases. Interestingly, *Granulicatella adiacens, Prevotella melaninogenica* and *Veillonella parvula* (all within the ten most abundant in the core microbiome) significantly differentiated the deceased patients. At this point it is not possible to conclude whether they play a role in the progression to fatal cases or are present in all groups because they are resilient to microenvironment changes.

Microbiota form complex ecosystems on human surfaces reflecting strong positive or negative interactions and studies on these communities should not be limited to report differences in presence or abundance. Networks of co-abundance or co-dependency are necessary to better understand the role of microbiota in health and disease (42). Accordingly, we analyzed the interaction between members of the bacterial community in each clinical group and found marked differences. Each network was composed of subclusters where its members had strong positive interaction between them. The composition of these subclusters varied in each clinical condition. A strongly interactive subcluster with *Prevotella, Veillonella* and *Actinomyces* species was present in the asymptomatic group and this subcluster partially remained in the ambulatory patients. Subclusters in the severe patients were completely different with *Lachnospiraceae, Gemella* and *Parvimonas* in a subcluster of the network of hospitalized patients and *Bacteroides, Rothia* and *Saccharibacteria* or *Selenomonas* and *Neisseria* in the network of the deceased patients. Thus, major changes occurred in the structure of bacterial communities as the severity of the disease increased. The possible role of these large changes in the pathogenesis and severity of COVID-19 remains to be studied. Perhaps these changes affect the nature of the local and systemic inflammatory response.

The way the members of these networks interact was yet another variable, and the network in the asymptomatic individuals was the more interactive community, with the larger number of links and a balance in negative and positive interaction (52%+ and 48%, respectively). This complex community, balanced in its interactions, could be considered a normal state or normonetting. The balance was gradually lost as severity of the disease increased and in ambulatory cases positive interactions increased to 73%, whereas in fatal cases most interactions were positive (91%). This altered networking or disnetting may result in a highly unregulated community. A previous study also found that the complexity of co-abundance networks was decreased in patients with severe COVID-19, indicating a reduction in the interaction between members of the bacterial community (6). A highly unregulated bacterial community may result in marked metabolic alterations and strong microenvironment changes. In fact, our metabolic deductions showed significantly large changes in metabolic pathways as the severity of disease increased (Figure 6). It was noticeable that the activity of most metabolic pathways was markedly and gradually increased from asymptomatic to ambulatory, to hospitalized and to deceased patients. This may suggest that the gradual looseness in negative interactions impacts metabolic activity of the bacterial community. The increased metabolites produced by the microbiota may have profound effects on the patient’s health (12).

COVID-19 is a complex multisystemic, multiorgan disease probably due to the ability of the virus to disseminate and invade several cell types of the body (1,2). Changes in the bacterial communities may also contribute to severity of the disease. Oral microbes and microbial molecules might directly enter the bloodstream and contribute to the pathogenesis of systemic diseases. Brain specimens and cerebrospinal fluid from individuals diagnosed with Alzheimer’s disease suggest that *P. gingivalis* could colonize the brain and induce neurodegeneration (43). Also, network analyses in oral microbiota have shown a significant correlation of disease-associated bacterial species with proinflammatory cytokines (44), and a significant correlation of the abundance of *Staphylococcus* in the nasopharynx with systemic levels of IL-6 and TNF has been reported (36). On the other hand, oral microbiota plays integral roles in maintaining host health systemically and locally. One example is the conversion by oral microbes of nitrate to nitrite, which is absorbed and converted to NO, important for the control of blood pressure and endothelial function (45). Thus, disruption of the oral microbiota may in several ways affect severity of COVID-19 disease.

In summary, we report significant changes in diversity, composition, and networking in the saliva microbiota of patients with COVID-19 and found patterns associated with severity of the disease. We report oral species associated with each clinical stage because of its presence or abundance, as well as infection with opportunistic pathogens. Patterns of networking were also found associated with severity of disease. A highly regulated community (normonetting) was found in healthy people whereas poorly regulated populations (disnetting) were characteristic of severe cases. Characterization of microbiota in saliva may offer important clues in the pathogenesis of COVID-19 and may also identify potential markers for prognosis of the disease.

## Financial support

This work was supported by CONACYT, Mexico (Call 2020-1, “Apoyo para Proyectos de Investigación Científica, Desarrollo Tecnológico e Innovación en Salud ante la contingencia por COVID-19, Grant 312992, to JT).

## Supplementary material

**Supplemental Table 1.**
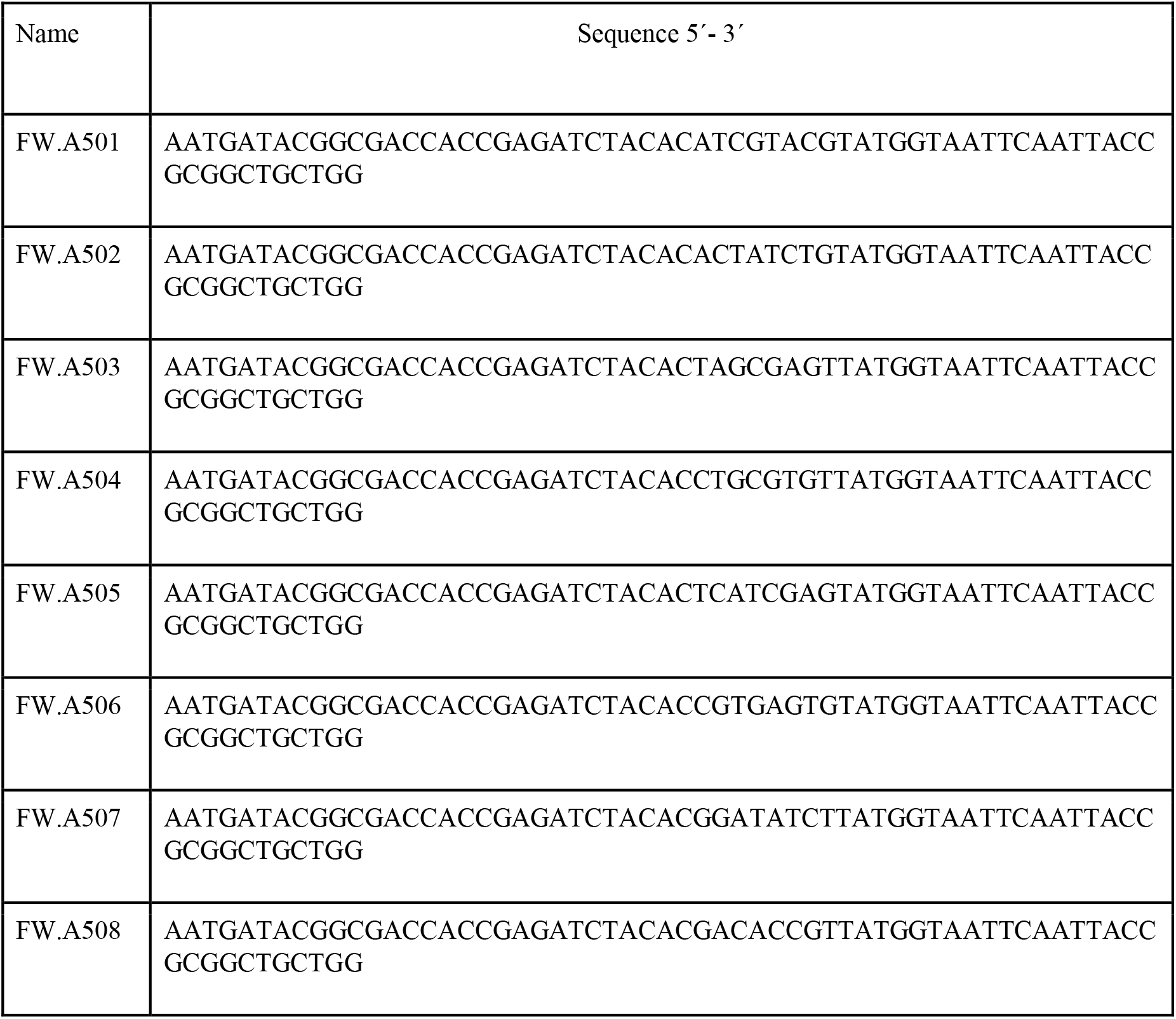
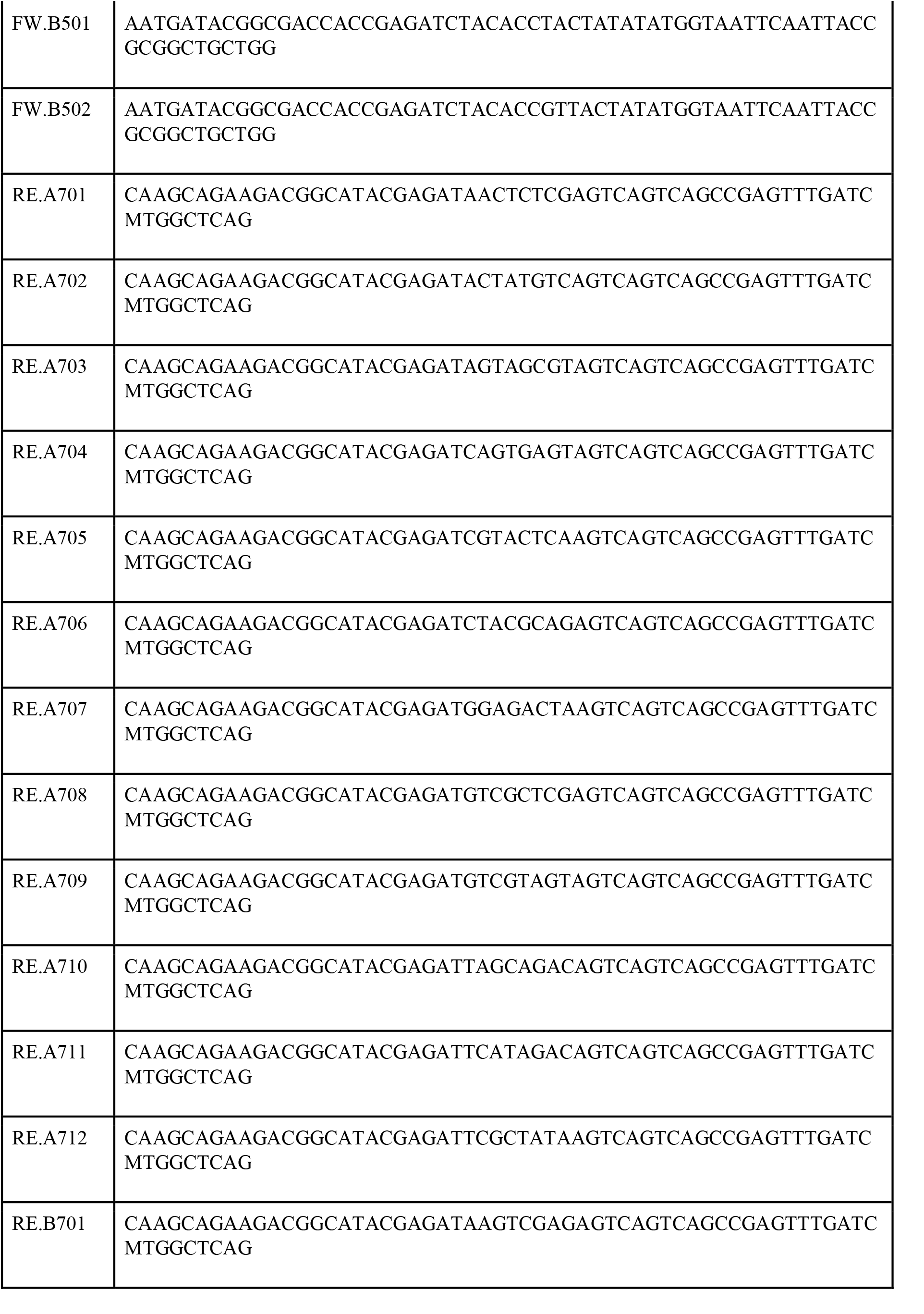
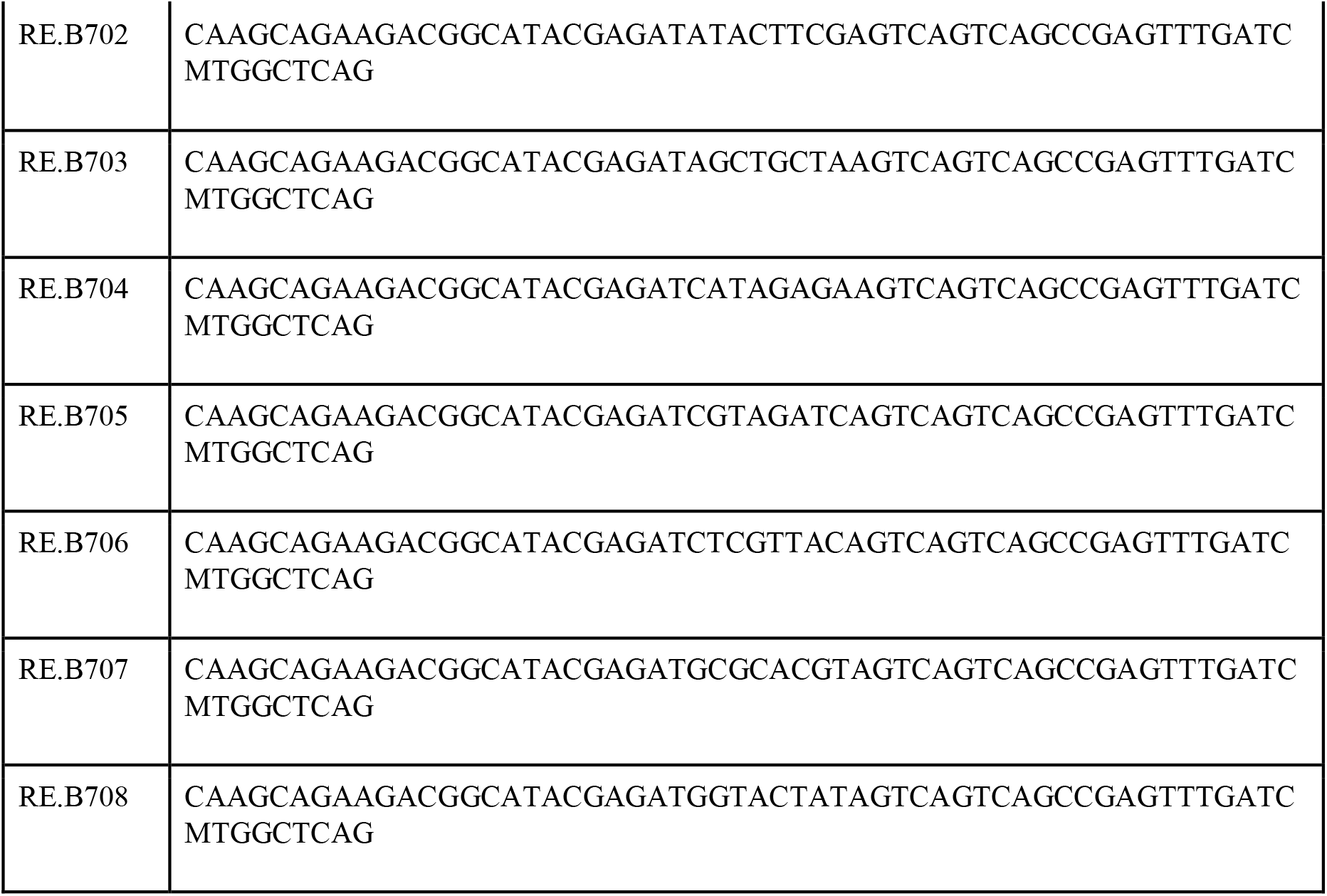
- List of primers used for the amplification of the hypervariable regions V1-V3 of the 16S gene

**Supplemental Table 2.**
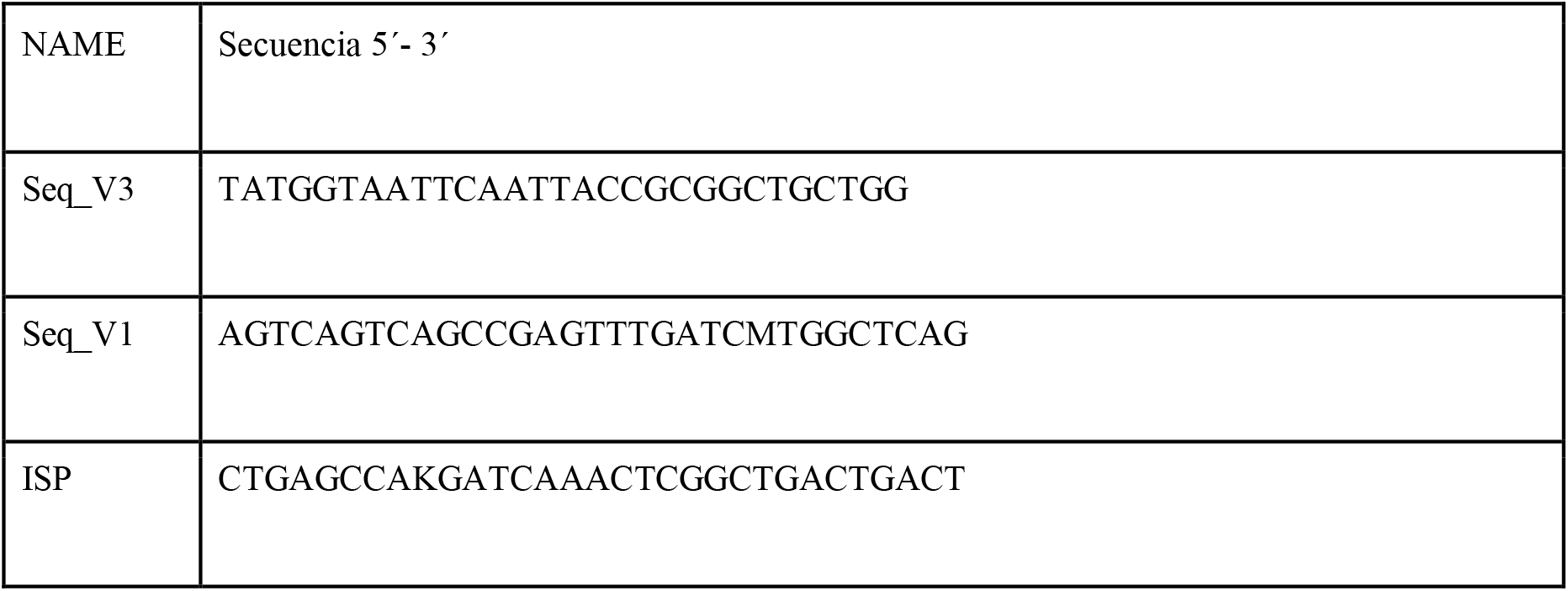
- List of primers used as adapters in the sequencing of the V1-V3 amplicons from the 16S gene, in the MiSeq instrument.

**Supplementary Figure 1.**
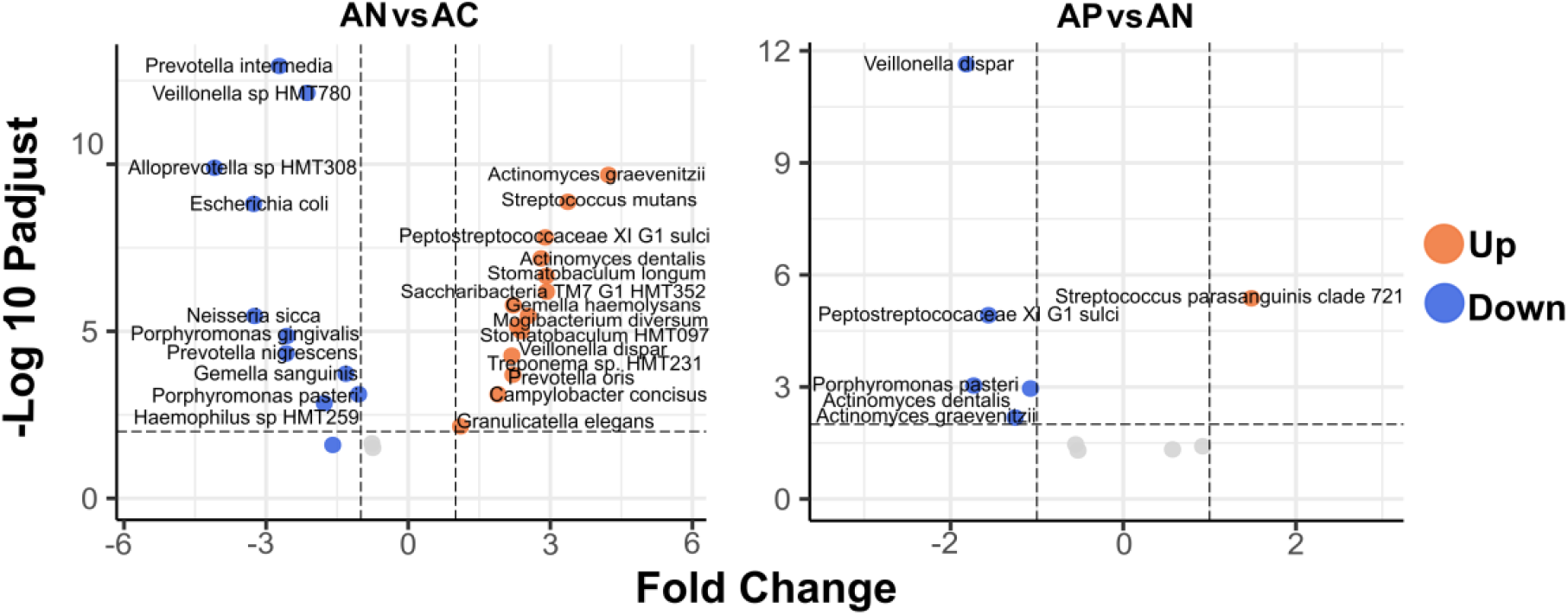
Pairwise comparison between clinical groups analyzed with the enhanced Volcano test. A) comparison between the ambulatory SARS-CoV-2 negative group and the asymptomatic control group; B) comparison between the ambulatory SARS-CoV-2 positive and the ambulatory SARS-CoV-2 negative. Species in orange dots presented increased abundance and those in blue decreased abundance.

**Supplementary Figure 2.**
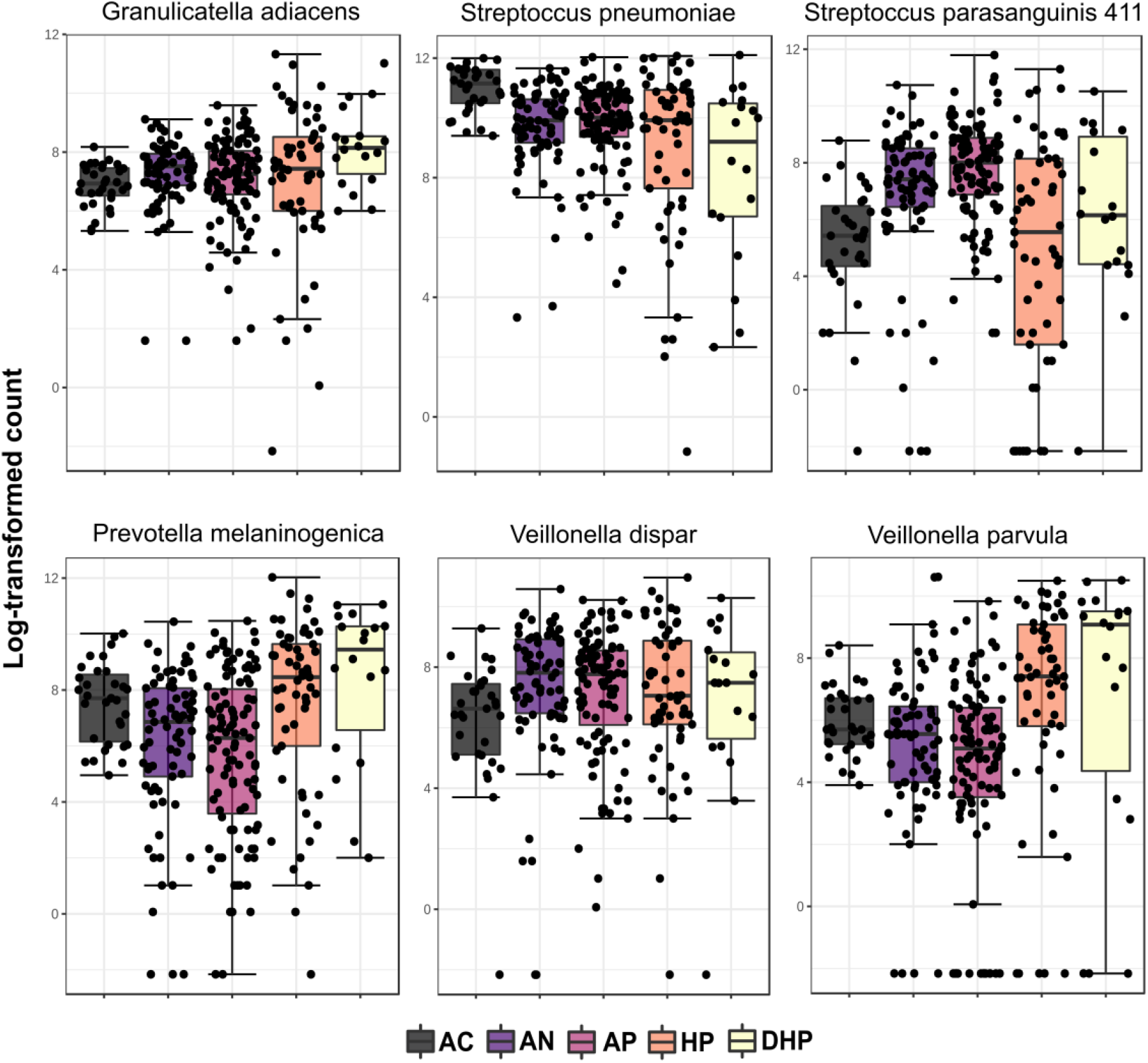
Boxplots of the relative abundance of species belonging to the core-microbiome, with over 1.0% abundance in at least 50% of all saliva samples in patients from all clinical groups.

